# Multidimensional analysis of drought response in an inter-specific tomato population (ToMAGIC)

**DOI:** 10.64898/2026.02.18.706544

**Authors:** Oussama Antar, Ana Rivera, Daniel Fenero, Lydia Serrano, Karen Alache, Aylin Kabaş, Jon Bančič, Mariola Plazas, Pietro Gramazio, Jaime Prohens, Santiago Vilanova, Joan Casals

**Author notes:** Corresponding author: Santiago Vilanova.

## Abstract

Drought stress poses a significant threat to agricultural productivity, particularly in regions with limited water availability. This study delves into the drought response in a multiparental interspecific tomato MAGIC population (ToMAGIC), developed by intercrossing *Solanum pimpinellifolium* (SP) and *S. lycopersicum var. cerasiforme* (SLC). A core collection of 139 recombinant lines, selected for their genetic diversity, was evaluated under both control and water stress conditions over two consecutive years. Phenotypic data were collected for 25 traits, including vegetative growth, flowering, fruit production, and physiological traits, providing a comprehensive assessment of drought response. Genome-wide association studies (GWAS) identified 15 significant genomic regions associated with drought response across eight chromosomes, highlighting key loci related to growth, earliness, fruit set, and physiological traits such as stomatal conductance and proline accumulation. Transgressive lines, such as S5_T_600 and S5_T_601, which exhibit enhanced drought resilience compared to the parental lines, were identified through genomic assisted selection, highlighting their potential as valuable breeding materials. The study emphasizes the importance of the ToMAGIC population in uncovering the polygenic nature of drought response. These findings offer valuable insights for developing drought-resilient tomato cultivars supporting agricultural sustainability in water-limited environments.

## 1. Introduction

Multiple abiotic stresses, such as heat, flooding and drought, increased markedly due to global climate change (IPCC, 2022; Yang et al., 2023). Among these stresses, drought stress represents a major threat to food security, impacting a large proportion of the global population, particularly those in arid and semi-arid regions (Raza et al., 2023). Water stress (WS) inhibits plant development by reducing the plant relative water content and water potential, leading to stomatal closure. As a result, plants suffer osmotic stress, restricted nutrient uptake, reduced photosynthetic activity, oxidative stress and inhibited growth (Nour et al., 2024).

Plants typically exhibit four types of response mechanisms to drought: avoidance, escape, tolerance or recovery (Fang & Xiong, 2015). Drought avoidance refers to the maintenance of physiological processes, including stomatal regulation, root system development, and other functions, during periods of WS. Drought escape is the ability to modify the life cycle, such as adopting a shorter life cycle, to evade the stress. Drought tolerance refers to a plant’s ability to cope with dehydration by employing various physiological processes, such as osmotic adjustment through the accumulation of osmoprotectants, soluble sugar content and peroxidase or superoxide dismutase activity (Ilyas et al., 2021). Drought recovery is the capacity of a plant to resume growth after experiencing stress (Zheng et al., 2023).

Tomato (*Solanum lycopersicum*), one of the most important and cultivated crops globally, is regarded as highly sensitive to abiotic stresses. Drought leads to reduced seed development and germination, decreased vegetative growth, and impaired reproductive processes, with the crop being particularly sensitive, especially during flowering and fruit enlargement stages (Xie et al., 2023).

Tomato relatives, such as *S. pimpinellifolium* (SP) and *S. lycopersicum* var. *cerasiforme* (SLC), are recognized as important sources for improving drought resilience (Albert et al., 2016; Martínez-Cuenca et al., 2020; Nakazato et al., 2008). SP and SLC were used for several years to introgress traits of interest into tomato cultivars through biparental crosses. However, biparental populations have limitations, such as low mapping resolution, which can hinder the precise identification of quantitative trait loci (QTLs) associated with complex traits. In contrast, multiparental advanced generation intercross (MAGIC) populations are powerful tools for QTL detection and have the added advantage of being easily incorporated into breeding programs(Arrones et al., 2020). Furthermore, combining MAGIC populations with approaches such as genome-wide association studies (GWAS) and genomic selection may uncover valuable information and generate new resources. Diouf et al. (2018) evaluated the genotype by environment (GxE) interaction under water deficit conditions in a tomato MAGIC population derived from four SLC and four *S. lycopersicum* var. *lycopersicum* (SLL) developed by Pascual et al. (2015). Plants were subjected to 25% irrigation reduction at the first flowering truss and by 50% at the second flowering truss, resulting in the identification of three QTLs regulated under water deficiency (*RIP9.1*, *RIP10.1* and *SSC1.2*).

In this study, an interspecific tomato MAGIC population (ToMAGIC) was assessed for drought response. The population was previously developed through intercrossing four SP and four SLC accessions using a funnel scheme including three cycles of intercrossing followed by five generations of selfing and has undergone extensive recombination events, offering high potential for identifying valuable traits. Each founder is estimated to contribute, on average, approximately 12.5% to the overall population (Arrones et al., 2024; Gramazio et al., 2020). Previously, the eight founders of the ToMAGIC population were assessed for drought deficiency in four different conditions (in greenhouse, as plantlets, in pots under both short and long cycles, and in the open field). The founders displayed a large phenotypic variation in response to water deficit when subjected to a 50% reduction in water supply (Antar et al., 2025).

The drought stress in the ToMAGIC population with 139 recombinant lines was studied over two years. During this period, 25 vegetative, earliness, flower, fruit, agronomic, and physiological traits were analyzed. This allowed the identification of patterns of phenotypic variation associated with control and water stress (WS) conditions. Genome-wide association studies (GWAS) were conducted to identify genomic regions associated with the drought response. Genomic assisted selection was applied to select superior transgressive lines under both stress and control conditions.

## 2. Materials and methods

### 2.1 Plant materials and growing conditions

A core collection of 139 recombinant lines from the ToMAGIC population (CCToMAGIC), selected based on hierarchical clustering of phenotypic data from the 354 recombinant lines, representing the phenotypic and genetic diversity of the whole population, was screened for drought response. The eight founders of the ToMAGIC population, which included four SP accessions (SP1, BGV007145; SP2, BGV006454; SP3, BGV015382; SP4, BGV013720) and four SLC accessions (SLC1, BGV007931; SLC2, LA2251; SLC3, PI487625; SLC4, BGV006769) were also included in the screening, except SP1, which failed to germinate.

Plants of the CCToMAGIC and their parentals were cultivated in an unheated glasshouse at the Agròpolis-Parc UPC experimental station (Viladecans, Spain; UTM: 41°17’21.5” N 2°02’42.5” E) from March to June over two consecutive years (2023 and 2024). Each year, seedlings with two true leaves were transplanted into 18 L plastic container pots filled with a growing substrate (J-2 Universal Substrate, Burés SAU, Sant Boi de Llobregat, Spain; 50% v/v), peat moss (Green Terra Profesional, Green Terra, Riga, Letonia; 25% v/v), and perlite (Burés SAU, Sant Boi de Llobregat, Spain; 25% v/v). Plants were grown for 100 days after transplanting (DAT), with side shoots being removed weekly upon appearance. Two watering regimes were applied: control (C) and water stress (WS). Water stress was induced by withholding irrigation during two periods, respectively lasting 24 days (early period, WS1; 26-50 DAT in both 2023 and 2024) and 16 days (late period, WS2; 68-84 DAT in both 2023 and 2024) (Figure 1a), until plants exhibited severe wilting symptoms. After each WS period, pots were rehydrated to full water capacity (recovery period), defined as the soil moisture level of the control treatment, by watering until drainage occurred. Substrate moisture levels were continuously monitored using Watermark Watchdog sensors (Copersa, Vilassar de Dalt, Spain), with hourly data recorded from two pots per treatment. Control plants were irrigated optimally to maintain substrate moisture within 25–40% of volumetric water content, corresponding to field capacity, with excess water allowed to drain freely from the pots. A fertigation solution (Hakaphos Basaplant Green, Compo Expert, Barcelona, Spain; 1 g/L) was applied twice per week, except during WS periods, when fertigation was suspended to maintain consistent nutrient supply across treatments. Two climatic sensors (temperature and relative humidity) were installed at a height of 1 m within the canopy, recording data every minute and averaged hourly throughout the experiment. The temperature and vapor pressure deficit (VPD) were similar in both years, averaging 22.2°C and 1.11 kPa for 2023, and 21.1°C and 0.99 kPa for 2024 (Figure 1b).

**Figure 1.**
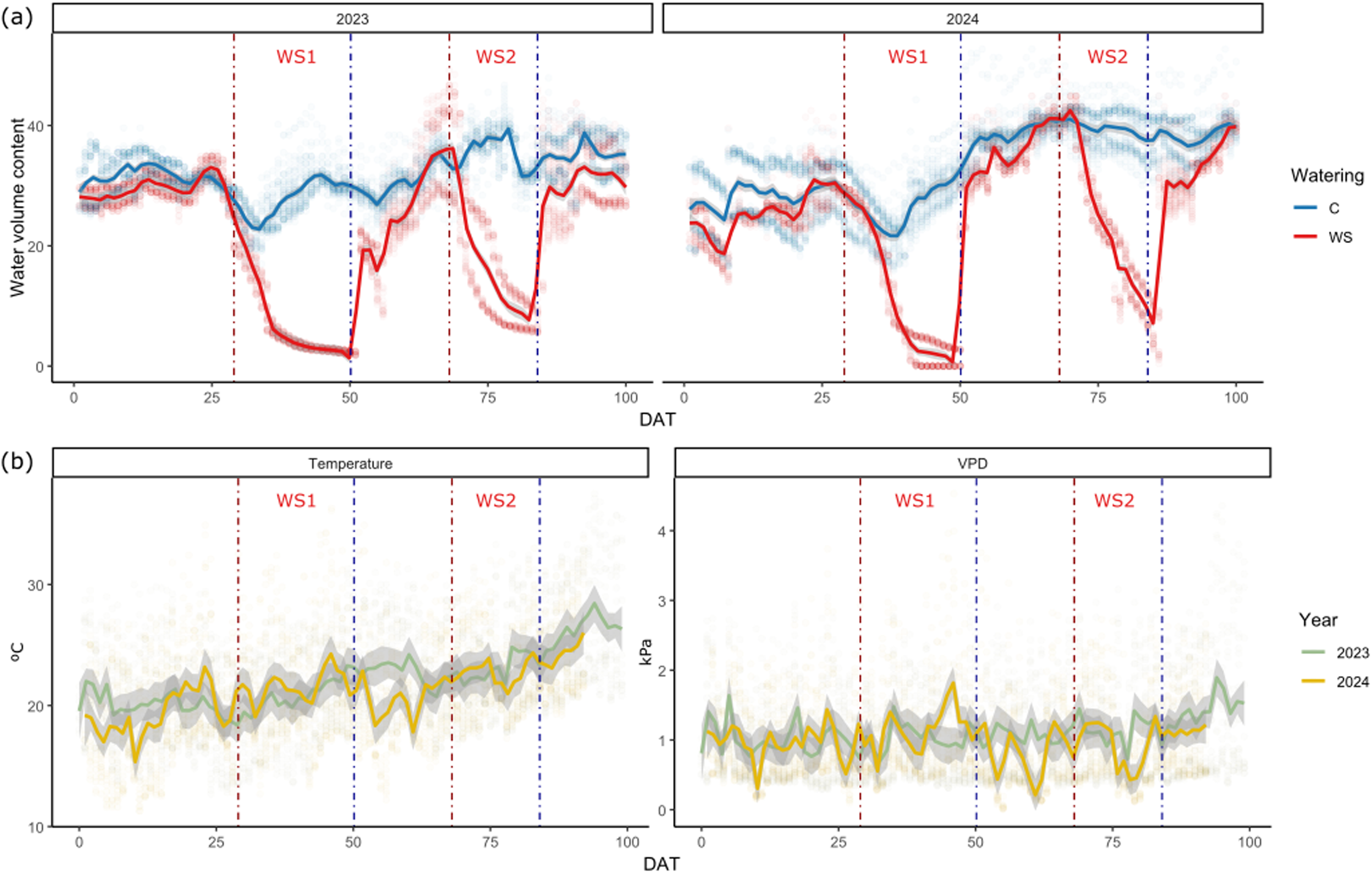
Experimental conditions during the 100-day crop cycle after transplanting (DAT, days after transplant). (a) Evolution of substrate water volume content in the control (C) and water stress (WS) treatments during the 2023 and 2024 experiments. (b) Comparison of temperature and vapor pressure deficit (VPD) between the two years. WS1 and WS2 indicate the two periods of water stress applied, while points in lighter shade represent hourly measurements taken throughout each day.

Each year, three plants per watering regime per CCToMAGIC parent or line were evaluated. Accessions were arranged in a three-block randomized design, with watering regimes applied in paired rows. The first and last rows were borders and were not part of the experiment. Bumblebees (*Bombus terrestris*) were introduced twice in each experiment, at the beginning and middle of flowering, to promote pollination.

### 2.2 Plant phenotyping

Twenty-five traits were phenotyped on an individual plant level throughout the crop cycle. Flowering earliness was assessed sequentially on the first eight trusses and expressed as the number of days from sowing (DAS) to the anthesis of the first flower in each inflorescence (FlowEarl1-8), as well as by the interval in days between two consecutive trusses (FlowTruss12-78, *e.g*. indicating interval length between trusses 1^st^ to 2^nd^ or 7^th^ to 8^th^). For trusses 1^st^ to 6^th^, the number of flowers (Nfl1-6) and fruits (Nfr1-6) was recorded to calculate fruit set (%), and the total number of trusses (TNTrusses) per plant was determined at the end of the experiment. Prior to the onset of the first WS period (22 DAT) and at the end of the WS period (49 DAT) leaf chlorophyll content (SPAD1 and SPAD2, respectively) (SPAD units) was measured on the first fully developed leaf in the upper part of the plant using a SPAD 502 meter (Konica Minolta, Tokyo, Japan). Stomatal conductance to water vapor (g_sw_, mol m⁻² s⁻¹) was measured at the end of each WS period only in the 2024 trial (g_sw_1 _and_ g_sw_2, respectively) with a LI-600PF porometer (LI-COR, Lincoln, USA). Leaf wilting was visually assessed at the end of both WS periods (wilting1 and wilting2) using a four-level ordinal scale: 1, less than 25% of leaves exhibiting wilting symptoms; 2, 25–50% of leaves affected; 3, 50–80% of leaves affected; and 4, severe wilting symptoms observed across the entire plant (Figure S1). Two representatives red-ripe fruits per plant from the 4^th^ or 5^th^ trusses were evaluated for fruit weight (FW, in g), soluble solids content (SSC, °Brix; PAL-1 Atago refractometer, Tokyo, Japan), and firmness (Durofel units: Agrosta Durofel, Compainville, France). To estimate final biomass production, senescent leaves were collected weekly, counted (LeafLoss.n) and weighed, then their cumulative weight was included in the final biomass estimates. At 90–95 DAT, plant height (PH, cm), stem diameter (measured at the base (DiamB) and above the third truss (Diam3), cm), and leaf length above the third truss (LeafL, cm) were measured. At the end of the crop cycle (100 DAT), all remaining fruits were harvested, ripe fruits were counted and weighed to determine total fruit number per plant as commercial yield (Yield.fw, g ripe fruit plant^-1^). To assess biomass partitioning, plants were separated into stems, leaves, and fruits, with fresh weight (fw) being determined immediately. The samples were then dried in a thermo-ventilated oven at 70°C for 72 h to determine dry weight (dw). Leaf biomass and stem biomass were combined and considered as VegetativeB.dw, while fruit biomass was expressed on both a fresh and dry weight basis (FruitB.fw, FuitB.dw). Total biomass was calculated as the sum of the three fractions (TotalB.dw). The harvest index (HI.dw, in %) was determined as the ratio of FruitB.dw to TotalB.dw (Table S1).

### 2.3 Proline extraction

Three samples per genotype per treatment were collected at the end of the first stress stage of the 2024 trial and immediately frozen at -20 °C. Proline content was then calculated following the method described byBates et al. (1973). Fresh leaf tissue (0.05 g) was extracted with sulfosalicylic acid (3%, w/v) and mixed with acid ninhydrin. The mixture was incubated in a water bath for 1 h at 98°C, cooled on ice for 10 min, and subsequently extracted with toluene. The absorbance of the organic phase was measured at 520 nm, and the concentration was quantified using an L-proline standard curve assayed in parallel. Results were expressed as µmol g⁻ ¹ dw.

### 2.4 Statistical analysis

Statistical analyses were conducted using RStudio software (version 4.4.1) (Racine, 2012). Raw data were filtered based on the Z-score method (Eq. 1). Points that had a Z-score above 3 were eliminated.

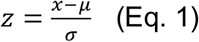

where x is the phenotypic data, µ and σ are respectively the mean and the standard deviation of each combination.

An analysis of variance (ANOVA) test was performed on the CCToMAGIC lines, where the total sum of squares was partitioned into components corresponding to genotype, treatment, their interaction, and residual error effects.

Furthermore, multiple Mixed Linear Models (MLM) were fitted for the analysis detailed below. The general form of the applied model (Eq. 2) was:

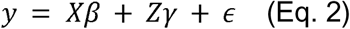

where y represents the observed response variable, *X* is the design matrix for the fixed effects β, *Z* the design matrix for the random effects γ, and ε is the residual error term. Specific models were tailored to the analysis goals as follows:

To identify traits with year-dependent (and potentially sign-reversing) water stress effects, we fitted a model including treatment, year, and their interaction as fixed effects, while genotype and the interaction between year and block as random effect using lme4 package(Bates et al., 2015).

Estimated marginal means (Emmeans) using “emmeans” package (Lenth, 2023) were calculated to evaluate the treatment effects across years, and contrast tests adjusted for Tukey method were conducted. Traits exhibiting a significant interaction between treatment and year were excluded from the analysis.

To assess the treatment effects at each stage for traits, such as earliness (FlowEarl, FlowTruss), Nfl, Nfr, and fruit set, wilting and g_sw_, which were measured sequentially at different stages, a model was conducted. The fixed effects consisted of treatment, stage, and the interaction between treatment and stage, while the random effects involve genotype, and the interaction between year and block. Emmeans were calculated, and a pairwise post hoc test adjusted using the Tukey method was applied.

#### 2.4.1 Drought indexes

Best Linear Unbiased Estimates (BLUEs) were calculated to perform multivariate analysis, compute drought indexes and perform GWAS. An MLM was employed using the “sommer” package (Covarrubias-Pazaran, 2016). In this model, the fixed effects consisted of genotype, treatment, and their interaction, while the random effects accounted for year and block.

Using the BLUEs extracted from the described above MLM, drought indexes were calculated as follows:

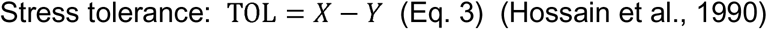

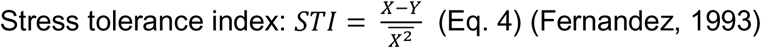

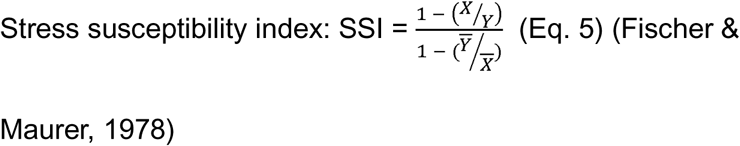

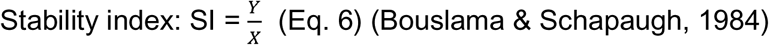

where *X* and *Y* are the BLUEs of a genotype under control and WS respectively, *X̄* and *Ȳ* are the population mean of the BLUEs under control and WS, respectively. *X̄*^2^ refers to the mean of the squared BLUEs under control conditions. All indices were calculated for all traits.

#### 2.4.2 Heritability and genomic assisted selection

Genomic Best Linear Unbiased Prediction (GBLUP) models were fitted for each treatment separately to estimate heritability and perform genomic-assisted selection. This model accounted solely for random effects including genotype, genomic relationship matrix (GRM) and the interaction between year and block using the “sommer” package. For each trait heritability (Eq. 7) was calculated using the Cullis method (Cullis et al., 2006), according to the following formula:

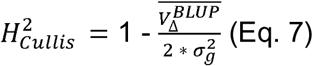

where σ_g_^2^ refers to genetic variance, and *V̄_Δ_^B̄L̄ŪP̄^* to the average standard error of the genotypic BLUPs.

Genetic estimated breeding values (GEBVs) extracted from the model were used to perform genomic assisted selection through a Smith-Hassel index (Eq. 8):

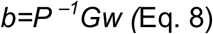

where *P* represents the phenotypic covariance matrix, *G* denotes the genetic covariance matrix, and *w* is the vector of weights assigned to each trait, with all traits receiving an equal weight of 1 (Hazel, 1943; Smith, 1936). The phenotypic covariance matrix was derived from the BLUEs, whereas the genetic covariance matrix was obtained from the GEBVs.

#### 2.4.3 Multivariate analysis

The BLUEs for traits significantly affected by treatment were standardized using Z-score normalization, followed by Principal Component Analysis (PCA). Boruta feature selection, an extension of the Random Forest method, was applied to identify the most relevant variables contributing to trait variation across treatments (Kursa & Rudnicki, 2010). Pairwise Pearson correlations were carried out between analysed parameters within each treatment (control and WS).

#### 2.4.4 Genome wide association studies (GWAS)

Genotypic data and a genomic relationship matrix (GRM) were combined with the drought indexes and the WS BLUEs to conduct a GWAS (Yu et al., 2005) for all traits from the “rrBLUP” package (Endelman, 2011).

A set of 6488 high-quality SNPs selected from an initial 4 268 587 variants, was obtained using a 12k probes tomato SPET panel, and previously filtered and retained by Arrones et al. (2024), who reported no evidence of genetic structure in the population. This set was used to perform the GWAS analysis using a mixed linear model (MLM) incorporating the GRM as a random effect. A genomic relationship matrix (GRM) (VanRaden, 2008) was calculated using “ASRgenomics” (Gezan et al., 2022).

Multiple testing was corrected with the Bonferroni threshold with a significance level of 0.05 (Holm, 1979). Associations were only considered significant if a SNP exceeded the Bonferroni threshold (5.11), defined as the −log10 of the desired overall alpha level (α = 0.05) divided by the total number of SNPs. Candidate genes were identified within the associated regions using the Sol Genomics Network database (Fernandez-Pozo et al., 2015).

## 3. Results

### 3.1 Phenotypic variation in the CCToMAGIC and its founders

A wide variation was found in the materials used. Table 1 displays the averages, ranges, and phenotypic variation observed for both the parental lines and the CCToMAGIC population under control and WS conditions, for the traits showing significant differences for genotype, treatment, or their interaction. Water stress induced a reduction of the values of several vegetative, fruit and flower-related traits and stomatal conductance (g_sw_), while resulting in an increase in traits such as proline (Table 1).

**Table 1:**
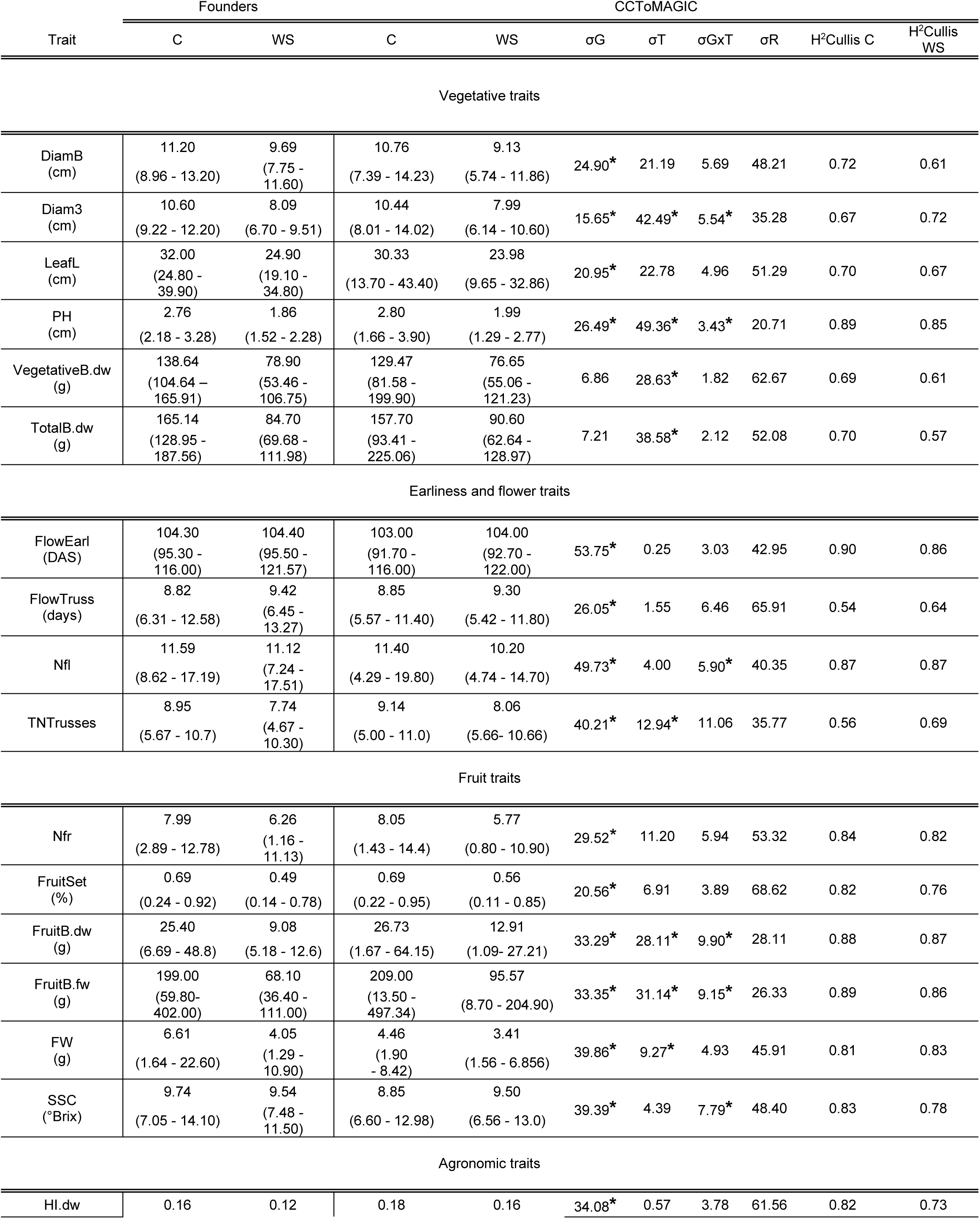
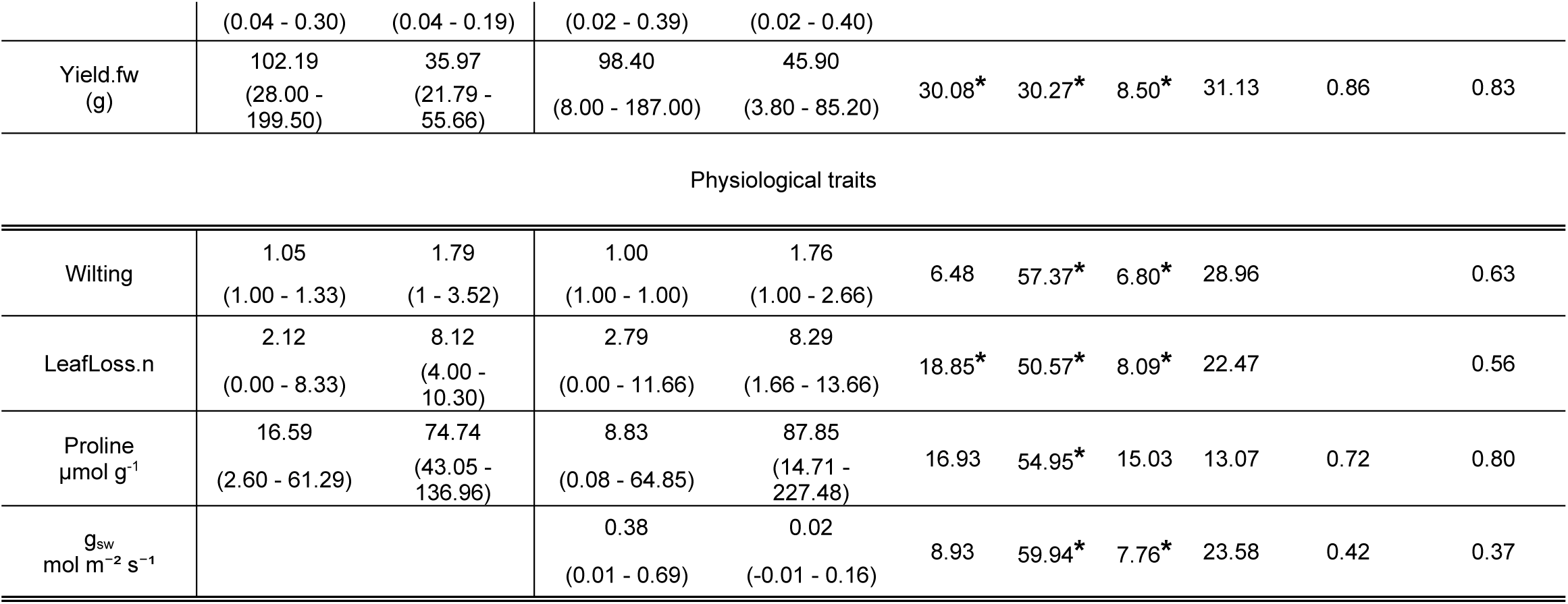
Means, minimum and maximum values of the CCToMAGIC and its founders, heritability (H^2^) under both water stress (WS) and control (C) conditions, percentage of total sums of squares explained by the G, T, and GxT, and residual effects (R) for the 22 significant traits for the genotype (G), treatment (T) and/or the GxT interaction. Sequential phenotypic traits were averaged. The presence of an asterisk (*) indicates that the factor has a statistically significant effect (p < 0.05).

Out of the 25 evaluated traits, significant effects (p < 0.05) were observed for 17 traits with respect to the genotype factor, 13 traits for the treatment factor, and 10 traits for the genotype × treatment interaction. Among the 22 traits with significant effects from the main factors and their interaction, the genotype’s contribution to the total sums of squares ranged from 6.48% (wilting) to 53.75% (FlowEarl). The treatment factor contributed between 0.25% (FlowEarl) and 59.94% (gsw). The interaction’s contribution was lower, ranging from 1.82% (Vegetative.B.dw) to 15.03% (proline). Heritability ranged in the control treatment from 0.42 (g_sw_) to 0.90 (FlowEarl) while in WS it ranged from 0.37 (g_sw_) to 0.87 (Nfl, and FruitB.dw) (Table 1).

The MAGIC population founders exhibited substantial phenotypic variation in both control and WS conditions. Both the CCToMAGIC and the founders exhibited comparable means under control and WS conditions. Proline was one of the traits that showed the largest variation in response to the treatment factor. The founders had mean values of 16.59 µmol g^-1^ under control and 74.74 µmol g^-1^ under WS, with ranges spanning from 2.60 µmol g^-1^ to 61.29 µmol g^-1^ in control and from 43.03 µmol g^-1^ to 136.96 µmol g^-1^ in WS treatment. In contrast, FlowEarl exhibited the lowest variation due to treatment. The founders presented mean values of 104.3 days in control and 104.4 days in WS, with ranges between 95.3 and 116 days under control and 95.5 to 121.57 days under WS.

High phenotypic variance was also observed in the CCToMAGIC population, with transgressive values recorded for most traits (Table 1). In CCToMAGIC, proline showed the greatest treatment-induced variation, exhibiting mean values of 8.83 µmol g^-1^ under control and 87.87 µmol g^-1^ under WS, with ranges from 0.08 to 64.85 in control and from 14.74 to 227.48 in WS. In contrast, FlowEarl displayed much lower variation. In the CCToMAGIC population, mean values were 103 days under control and 104 days under WS, with ranges of 91.7–116 and 92.7–122 days, respectively.

### 3.2 Trait variation across developmental stages

FlowEarl, Nfl, Nfr, fruit set, wilting and g_sw_ were measured sequentially. Traits related to earliness were assessed up to the 8^th^ truss, whereas flower and fruit-related traits were recorded up to the 6^th^ truss. Wilting and g_sw_ were measured twice, at the end of each WS period. A contrast test for sequential measurements analysis indicated that the treatment effect became significant from the 4^th^ inflorescence onward. FlowTruss showed a significant effect of WS starting from the interval between the 3^rd^ and 4^th^ truss. Wilting and g_sw_, were both significantly affected at both stages. Wilting increased slightly after the 2^nd^ WS period, whereas g_sw_ remained equally low in WS across both periods (Figure S2).

Water stress (WS) had a greater impact on flower and fruit-related traits (fruit set, Nfl, Nfr). Contrast tests revealed that fruit set and Nfr were significantly affected starting from the 2^nd^ truss onward. In the 1^st^ truss, the fruit set was low in both conditions compared to the later-developed trusses (Figure 2). Similarly to fruit set, WS started to affect significantly Nfr per truss from the 2^nd^ truss. The 4^th^ and 5^th^ trusses had the highest Nfr in both conditions, with 10.43 and 11.32 fruits in the control condition and 7.66 and 7.72 fruits in WS, respectively. Regarding Nfl, the difference starts to become slightly significant from the first truss. The highest number of flowers in control conditions was observed in the 5^th^ and 6^th^ trusses, while in WS conditions, the number of flowers was similar across the 3^rd^ to 6^th^ trusses.

**Figure 2.**
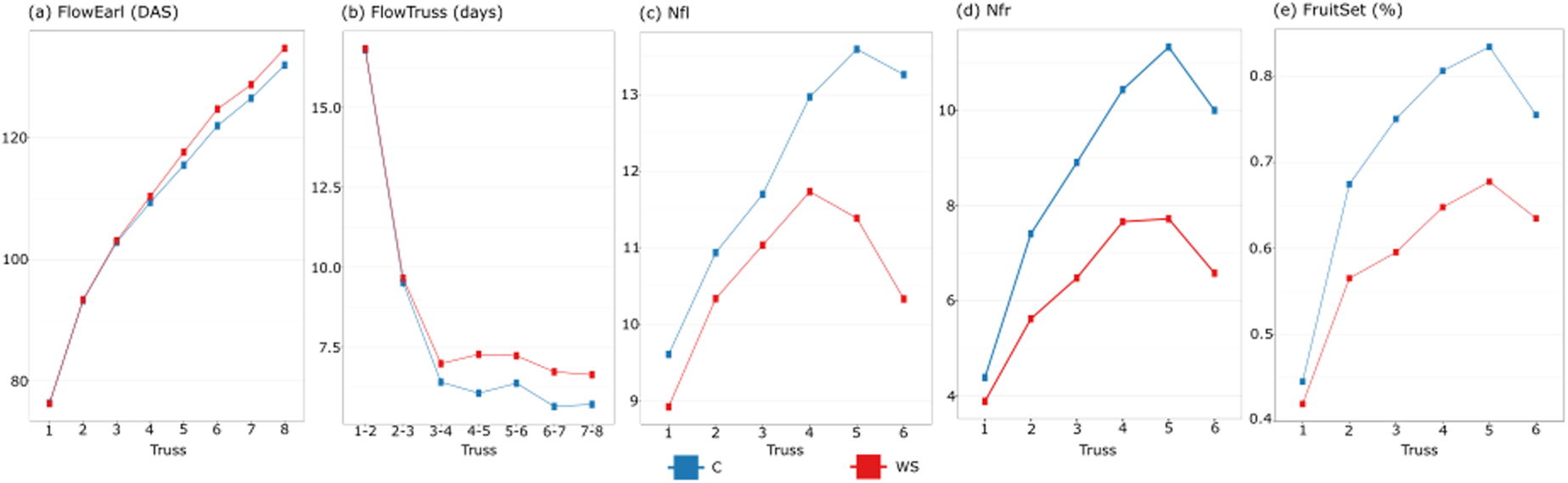
Sequential measurements of earliness and inflorescence traits. (a–b) Flower earliness (FlowEarl) and inter-truss interval duration (FlowTruss) in the first eight trusses; (c–d) number of flowers and fruits per inflorescence, and fruit set in the first six inflorescences. Plots show the estimated marginal means under water-stress (WS, red line) and control (C, blue line) conditions in the CCToMAGIC population.

### 3.3 Multivariate analysis and Boruta feature selection

A Principal Component Analysis (PCA) conducted with traits showing a significant effect for the treatment and/or the interaction revealed two distinct clusters based on treatment, indicating a significant difference for trait expression among the treatments (Figure 3). The WS condition showed a more compact distribution, whereas the control condition was more dispersed, indicating that a broader phenotypic spectrum was expressed under control than under WS. The first two principal components (PCs) accounted for 56.8% of the total variation. PC1, accounting for 44.7%, was positively associated with the separation between the two clusters corresponding to the different treatments. PC2, which explained 12.1% of the variation, can be positively associated with the genetic variance and other unexplained factors.

**Figure 3.**
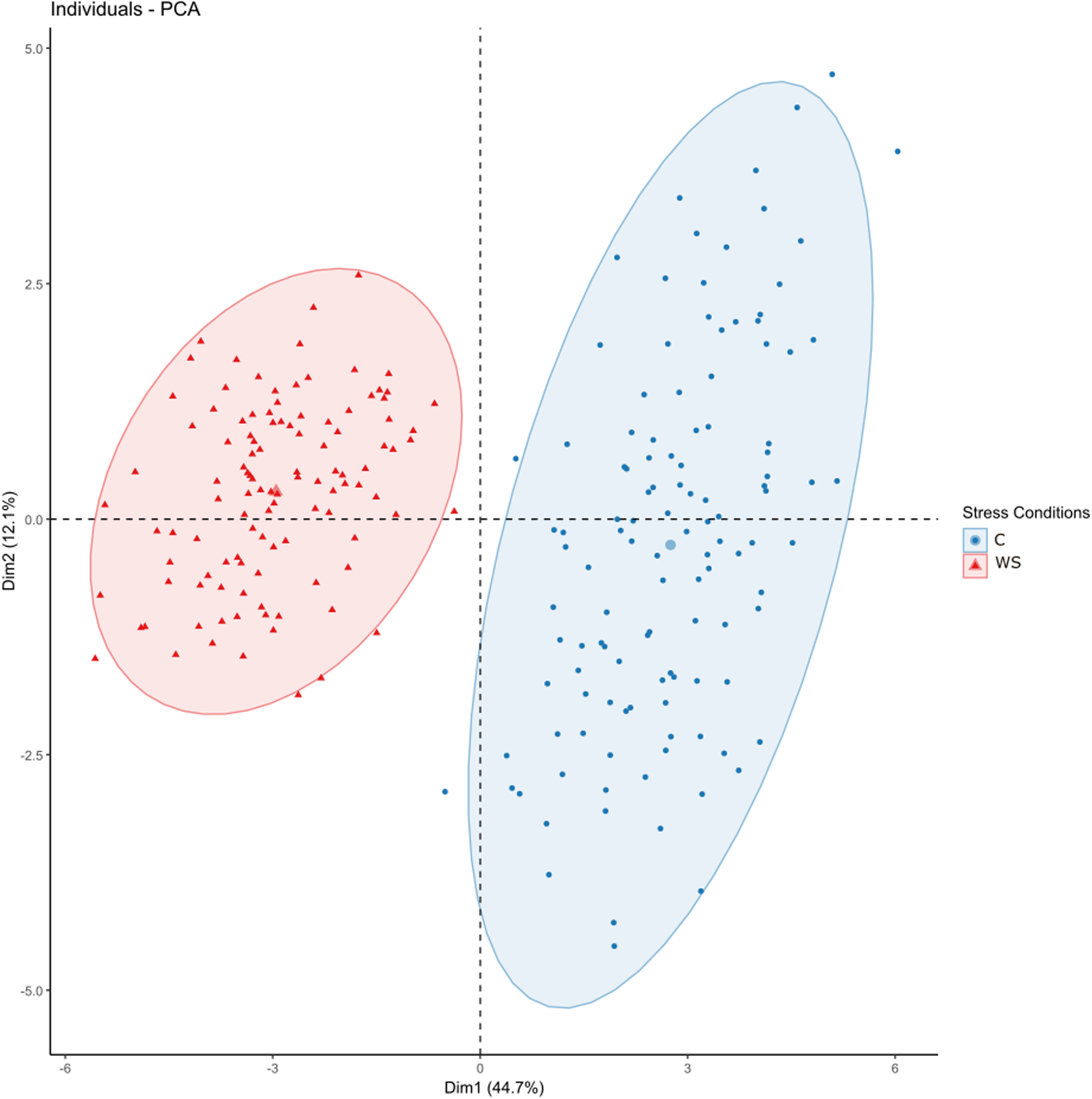
Principal component analysis (PCA) for the 22 traits showing a significant (p < 0.05) treatment and/or genotype × treatment interaction. PC1 (44.7%) account for the largest proportion of variation, driven by treatment differences. PC2 (12.1%) represents a smaller fraction of the variation, influenced mainly by genetic factor. Blue, control conditions; red, water stress treatment.

Boruta feature selection identified wilting, proline and g_sw_ after the two WS periods, and vegetative traits (VegetativeB.dw, TotalB.dw, DiamB, LeafL) as the best predictors of WS response (Figure 4). Additionally, LeafLoss.n, earliness (FlowEarl5, FlowTruss67, FlowTruss45 FlowTruss56). Fruit and yield related traits (FruitSet2, Nfr2, Yield.fw and fruitB.dw) were also recognized as key indicators of WS. Traits such as FlowTruss78, Nfl3, FlowEarl6, Nfl4, and Nfl5 were classified as tentative because they were at the threshold of significance.

**Figure 4.**
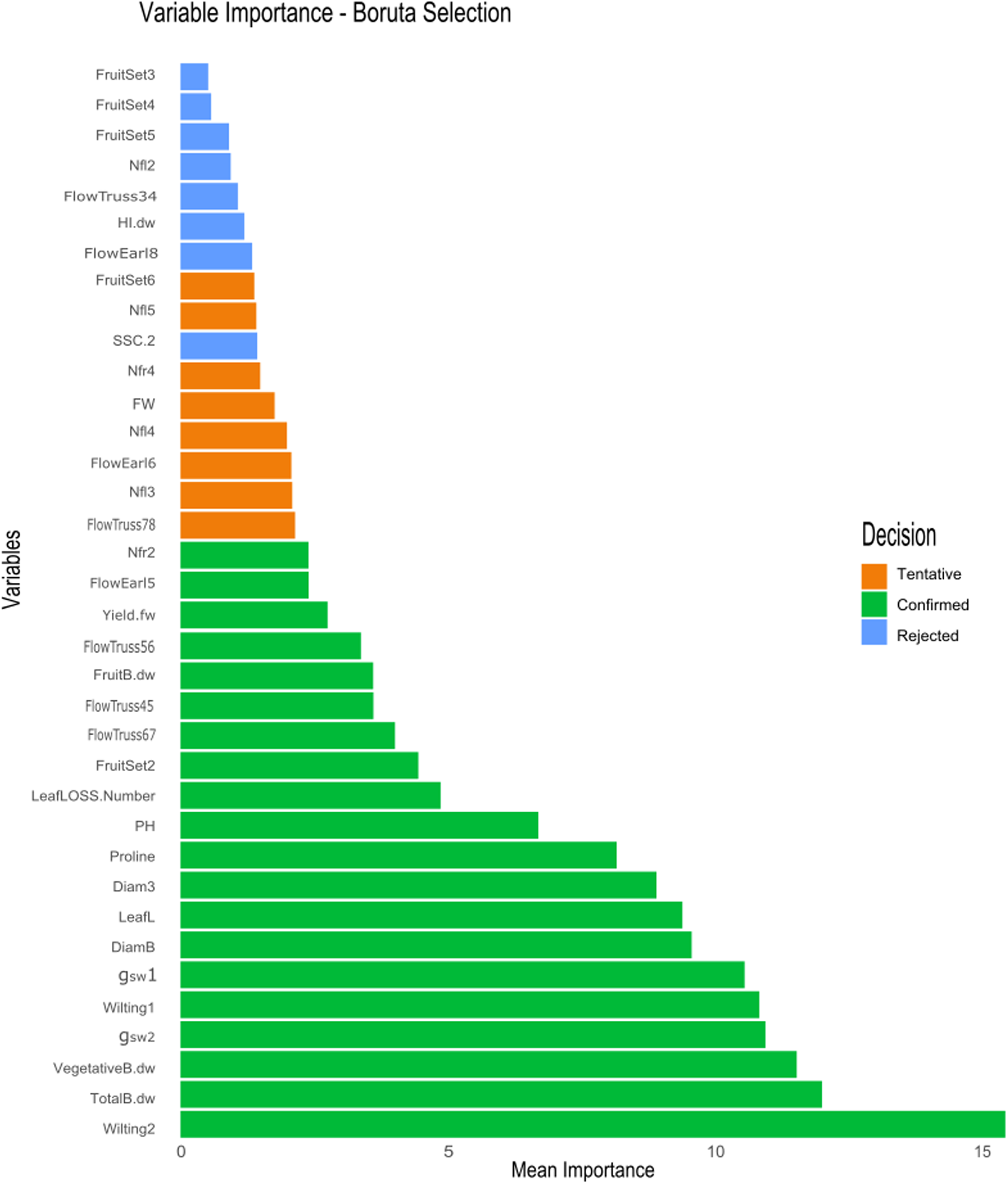
Boruta selection plot, where green bars represent significant traits associated with the treatment, orange bars represent uncertain traits, and blue bars represent traits that had an irrelevant association with the treatment. x axis represent mean importance.

The correlation analysis showed a diverse pattern of associations between traits in both conditions (Table S2). In control conditions, VegetativeB.dw was significantly correlated with TotalB.dw, PH, Diam3, FruitSet2, proline, and g_sw_2. In WS, TotalB.dw was significantly correlated with vegetative traits (VegetativeB.dw, PH, Diam3 and DiamB), leaf physiological traits (wilting, LeafLoss.n), fruit and agronomic traits (FruitB.dw, Yield.fw), SSC, and proline content. The correlations with vegetative traits, g_sw_2, FruitB.dw and Yield.fw traits were positive, while the correlation with proline, SSC and leaf physiological traits was negative. Proline content showed a significant negative correlation with g_sw_2 (r=-0.67, p < 0.0001), all vegetative and agronomic traits, but a significant positive correlation with FlowTruss45, leaf physiological traits and SSC (r=0.24, p = 0.003) (Table S3).

### 3.4 CCToMAGIC transgressive lines selection

Genomic assisted selection (GAS) was performed using the traits Wilting2, TotalB.dw, VegetativeB.dw, PH, FlowTruss45, and Yield.fw, Nfr4, FruitSet4. The first six were selected based on Boruta selection and agronomic relevance; variables from the 4^th^ truss (Nfr4 and FruitSet4) were added as representatives of the more affected truss in the sequential analysis (Figure 2d-e). Four lines (S5_T_504, S5_T_600, S5_T_601 and S5_T_724) were selected as the most tolerant and transgressive lines (Figure 5). Line S5_T_601 had the highest selection index under WS (5.47), followed by S5_T_600 (5.07), S5_T_504 (4.61) and S5_T_724 (4.46). In control conditions, the best performers were S5_T_505 (5.23) and the parental line SLC3 (5.18). In some cases, these selected lines also showed a good performance under control treatment, e.g. S5_T_600 (3.92 in control treatment, ranking sixth), S5_T_601 (3.21, 13^th^) and S5_T_504 (2.9, 14^th^). Parental lines, SP2 and SP4, had a selection index of 3.61 and 2.22 under WS, while SLC3 and SLC1 had a selection index of –4.81 and -4.39, respectively. In control conditions, SP3 had the lowest selection index (-3.01), followed by SLC1 and SP4 with a selection index of –2.72 and – 2.35, respectively.

**Figure 5.**
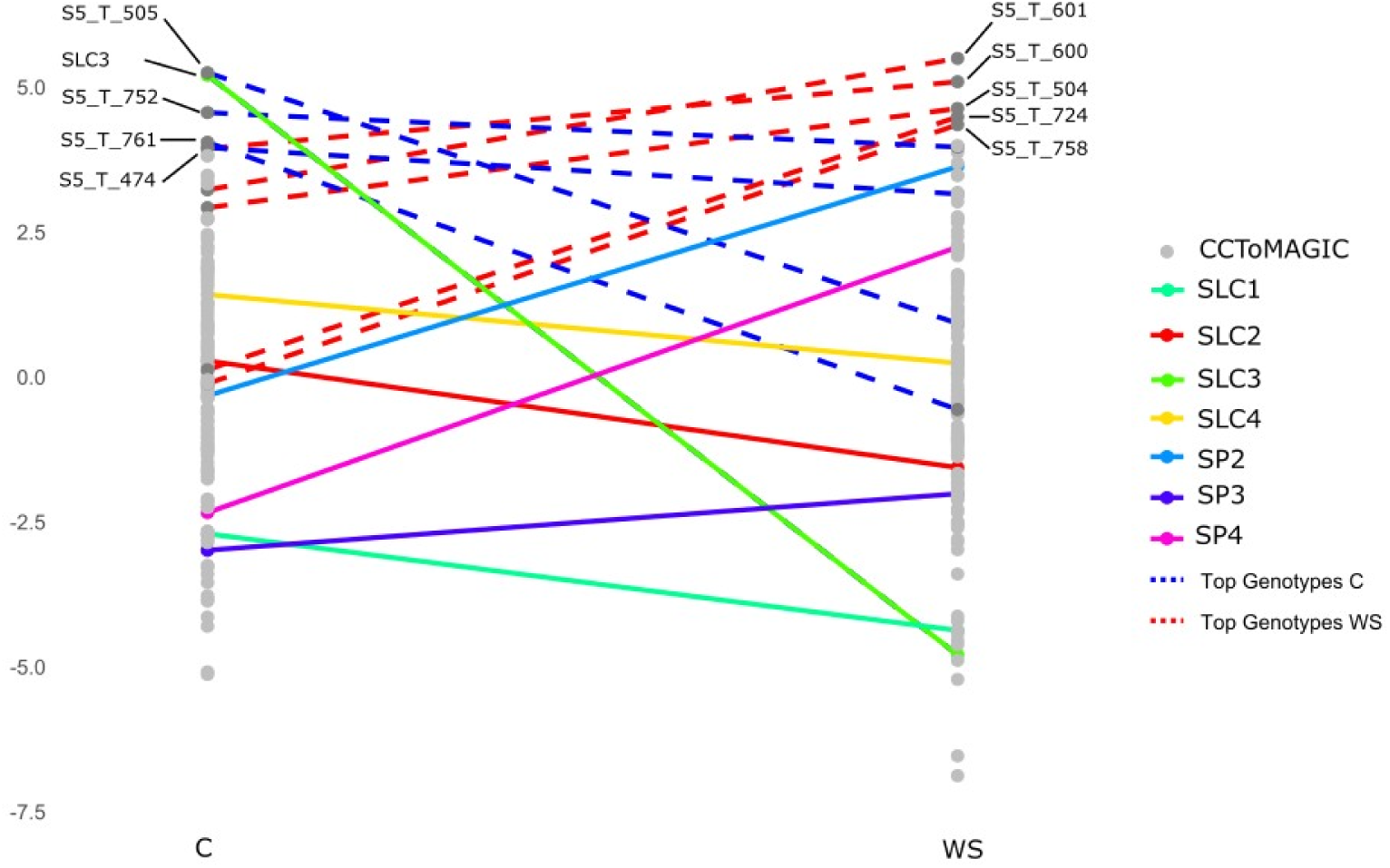
Selection index plot representing selection done based on seven traits in water stress (WS) and six in control conditions. Dots represent the CCToMAGIC and the founders. Coloured lines represent the founders, and black lines represent the top five genotypes with the highest selection indexes in control and WS treatments.

The S5_T_600 line ranked second in WS and sixth in control and experienced a 10.03% and 13.33% increase in WS compared to control regarding FruitSet4 and FlowTruss45 (Table 2). For Nfr4, S5_T_600 exhibited a 6.97% reduction under WS relative to C. Despite this, S5_T_600 belonged to the fourth quantile, exhibiting a high number of fruits in both conditions. Regarding TotalB.dw and Yield.fw, S5_T_600 suffered a 43.89% and 52.49% reduction in WS, respectively, but remained among the fourth quantile in terms of the most productive.

**Table 2.**
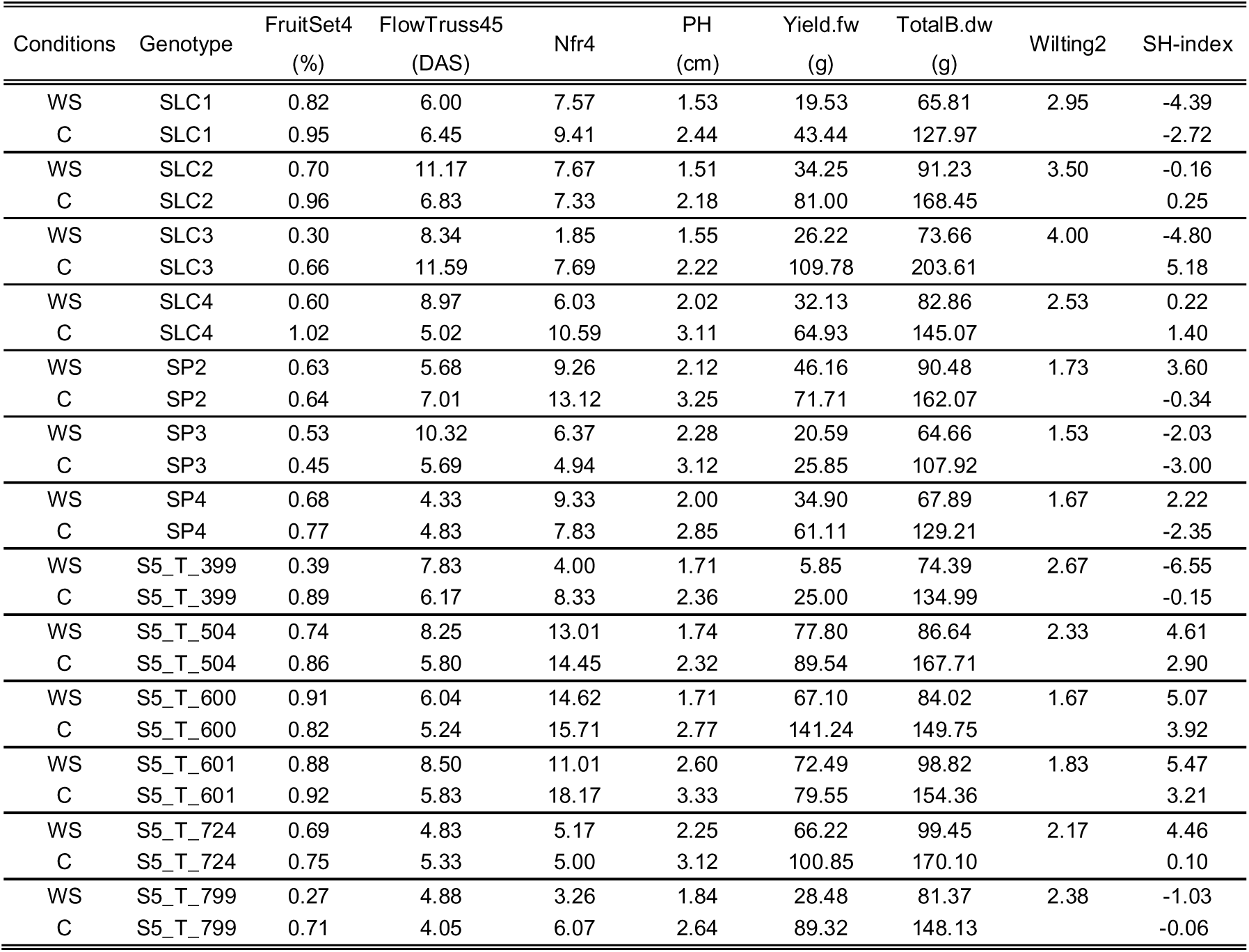
Trait values under control (C) and water stress (WS) conditions, and Smith–Hazel selection index (SH-index) of the parental lines, the four CCToMAGIC lines with the highest SH-index (S5_T_504, S5_T_600, S5_T_601, and S5_T_724), and the two lines with the lowest SH-index under water stress (S5_T_399 and S5_T_799).

Among the parental lines, SLC3, which had the lowest selection index, exhibited significant deviations in multiple traits under WS compared to control conditions. For FruitSet4, it was subject to a 55.02% reduction due to WS. Regarding FlowTruss45, SLC3 showed an increase of 28.02% due to WS. For Nfr4, WS drastically impacted this trait, leading to a 75.92% reduction, while it induced a 30% reduction in PH. For Yield.fw, and TotalB.dw, WS led to a severe reduction of 76.11%, and 63.82%, respectively (Table2).

### 3.5 Detection of genomic regions associated with drought response

Through GWAS, several genomic regions associated with drought response were identified, with SNPs surpassing the Bonferroni threshold value (5.11). GWAS analysis was performed with four indices and BLUEs in WS for each trait. A total of 22 significant SNPs were identified using SI, followed by 12 through BLUEs values in WS, 10 through TOL, 5 through SSI, and 2 through STI. Candidate genes were suggested based on their functions (Table S3).

Several SNPs associated with drought response were identified for vegetative traits, including PH, Diam3, TotalB.dw. For PH, a genomic region on chromosome 1 at 81.9 Mb was found to be associated with drought response. For TotalB.dw, an associated genomic region was identified on chromosome 3 at 56.2 Mb. For Diam3, a genomic region on chromosome 8 between 55.6 Mb and 56 Mb was identified. As a result, three genomic regions were identified as associated with these traits (Figure 6).

**Figure 6.**
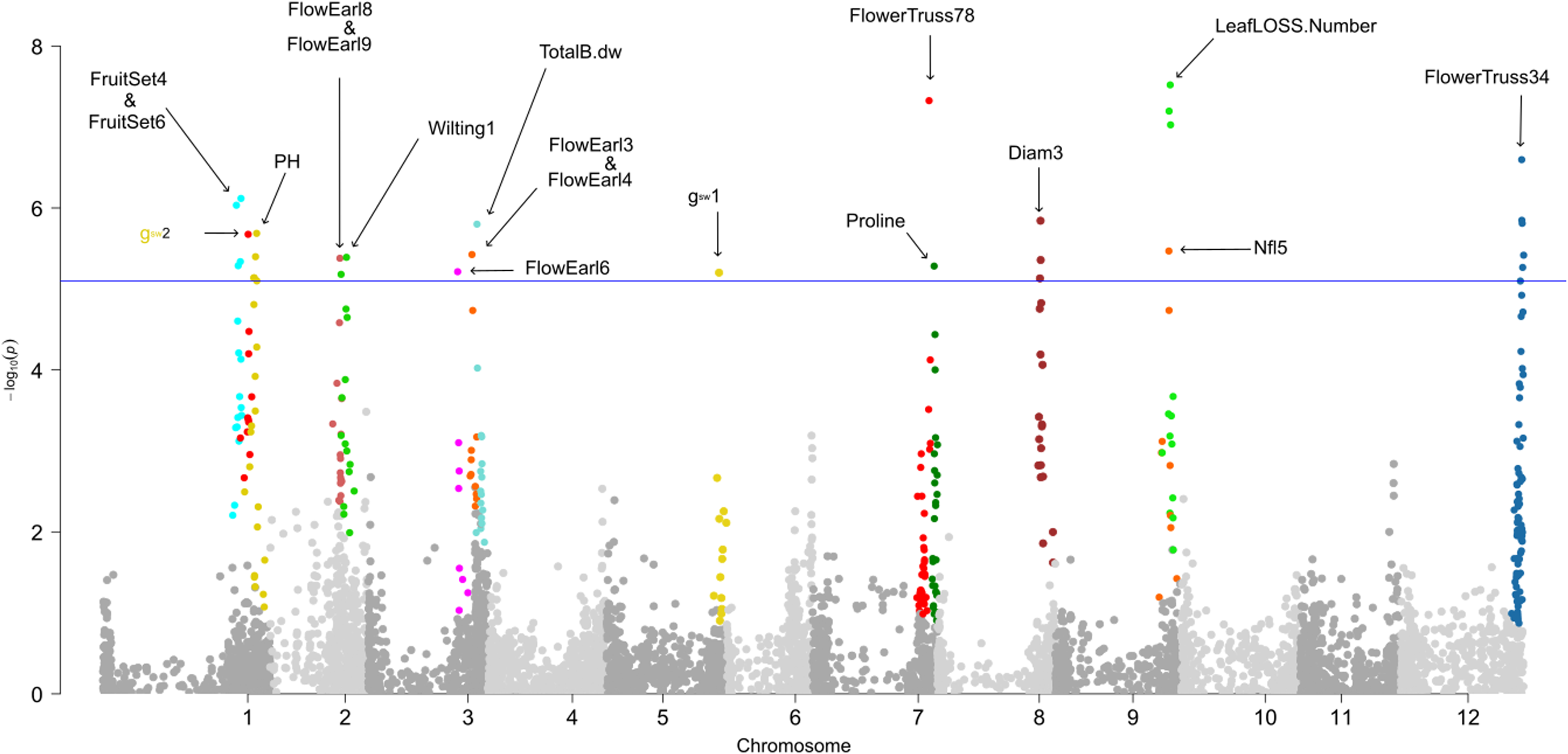
Manhattan plot of genome-wide association analysis (GWAS) of water stress response in the phenotypic spectrum and SNPs across 12 chromosomes using 6,488 SNPs. Horizontal line denotes the Bonferroni threshold -log10(p) value of 5.11. Significant associations exceeding this threshold are highlighted with coloured dots.

Several genomic regions were identified as associated with drought response in earliness-related traits. On chromosome 3, a region at 58.96 Mb was found to be associated with FlowEarl3 and FlowEarl4, while another at 48.48 Mb was associated with FlowEarl6. On chromosome 2, a SNP at 37.9 Mb was associated with later-stage earliness traits FlowEarl8 and FlowEarl9. For the day interval between two consecutive trusses, a SNP at 63.3 Mb on chromosome 7 was detected as being associated with FlowTruss78. Regarding FlowTruss34, the genomic region on chromosome 12 between 65.6 Mb and 65.68 Mb was identified as being associated with drought response. For drought response related to fruit and inflorescence traits, an association with Nfl5 was identified on chromosome 9 at 61.4 Mb. Additionally, two genomic regions associated with drought affecting FruitSet4 and FruitSet6 were identified on chromosome 1 in a genomic region between 72.49 Mb and 74.97 Mb (Figure 6).

For proline accumulation after the first WS period, a SNP on chromosome 7 at 66 Mb was found to be significantly associated. In terms of Stomatal conductance of water (g_sw_), which was measured twice, each time after a drought period. Two genomic associations were detected, one on chromosome 5 at 61 Mb after the first drought period and another on chromosome 1 at 78.8 Mb after the second drought period. Wilting and leaf loss are drought stress indicators. As a result, two genomic regions were identified as being associated with wilting after the first drought period (wilting1) and total leaf loss number (LeafLoss.n). A SNP on chromosome 2 at 41.4 Mb was found to be associated with Wilting1, while on chromosome 9, a genomic region was associated with LeafLoss.n between 61.7 Mb and 64.5 Mb (Figure 6).

## 4. Discussion

Drought response is a quantitative trait characterized by complex phenotypic expression and genetic regulation, whereby many genes are involved with minor effect(Chloupek et al., 2010). Given the complexity of the genetic control of drought response, marker-assisted selection has not contributed significantly to the improvement of cultivars in dry environments (Khatun et al., 2021). In this study, a multidimensional genotypic and phenotypic evaluation was performed using a core collection of 139 lines from the ToMAGIC population, along with their genetically diverse founders (SLC and SP). This material was assessed for drought response for two consecutive years, applying two water regimes (control and WS conditions). The founder parentals originate from various climatic conditions, ranging from humid and hot regions to arid deserts, contributing to the population’s broad genetic variation (Gramazio et al., 2020; Martínez-Cuenca et al., 2020). Beyond the inherited genetic diversity and ecological adaptations, during ToMAGIC development, intensive recombination events occurred, leading to a population with high genetic reshuffling(Arrones et al., 2024). The multidimensional assessment included phenotypic analyses describing factor variances, effects, heritability, and correlations. Sequential measurement analysis identified the developmental stages most sensitive to drought stress for earliness, Nfr, Nfl, fruit set, wilting, and g_sw_ sequence traits. GWAS revealed genomic regions associated with drought–trait interactions, with distinct associations observed at different developmental stages for the same trait. Finally, genomic assisted selection revealed transgressive lines exhibiting enhanced drought resilience.

Drought stress affects all stages of tomato plant development, leading to substantial reductions in fruit yield, flower production, and various morphological changes. However, there is no optimal single method for drought response evaluation, and the choice of assessment should be based on the plant material and experimental conditions (Flores-Saavedra et al., 2023). Regarding earliness traits, previous studies reported that plants experiencing drought stress may adopt an escape mechanism by flowering earlier (Verslues et al., 2023). However, in our findings, WS led to an overall delay in flowering across the population, with significant WS effects on earliness becoming evident from the fourth inflorescence onward. Among the parental lines, it was observed that founders originating from arid regions and low precipitation (SLC1, SP4) exhibit an earlier flowering in both conditions without any effect for WS application, which suggests an adaptation for drought escape due to their native environment (low annual mean precipitation). Conversely, other founders displayed varied responses. While SP2 and SLC3 experienced an earlier flowering under WS, SLC2, SLC4 and SP3 experienced a delay in flowering when subjected to WS (Table 2).

Fruit set ratio has proven to be a reliable trait for distinguishing between drought-tolerant and sensitive genotypes. Drought stress affects pollen/anthers development and causes irreversible growth abortion of anthers at the tetrad and uninucleate microspore stages (Lamin-Samu et al., 2021). Pollen fertility is reduced due to an alteration in sucrose and starch levels induced by drought stress(Hu et al., 2020). In our study, the greatest fruit-set reduction (∼20 %) occurred from the 3^rd^ truss onward (notably the 3rd, 4th, and 6th), but this mainly reflects the timing of stress imposition, as the most affected inflorescences depend on when WS is applied. Because WS experiments typically begin around the emergence of the 1^st^ inflorescence, our results can be extrapolated to similar experimental designs. Consistent with Grozeva and Ganeva (2024), we also found that WS reduced fruit number more strongly than flower number.

Genomic selection has been shown to outperform marker-assisted selection, providing higher prediction efficiency for abiotic stress tolerance (Cerrudo et al., 2018). Its effectiveness was demonstrated in major crops such as wheat, maize, and rice for both drought and heat tolerance (Budhlakoti et al., 2022). In maize, Das et al. (2020) reported genetic gains of 0.110 and 0.135 t/ha/yr for grain yield under drought conditions, and 0.038 and 0.113 t/ha/yr under waterlogging across two populations. In our study, traits used in the genomic assisted selection were chosen for being WS indicators, traits highly affected by drought and for their commercial value. Parental lines such as SP2 (coming from a coastal zone with low precipitation (49 mm annual mean precipitation) and high temperature (24°C mean annual temperature) (Martínez-Cuenca et al., 2020) was identified as the best parental line in WS, while in control conditions it had a low SH index. Our genomic assisted selection findings align with a previous study where SP2 was selected as the most tolerant line among the eight founders. On the other hand, SLC3, coming from a tropical climate with high annual precipitation (2274 mm) and moderate mean annual temperature (18°C), was identified as the most affected parental line in WS, while in the control condition it was the best performer with the highest yield and biomass production. Therefore, three CCToMAGIC transgressive lines (i.e., S5_T_600, S5_T_601 and S5_T_504) were found to outperform SP2 in WS and maintain a high selection index also in control conditions, shedding light on the importance of this population, especially by combining desired traits from all parental lines.

Plant interaction with drought is a polygenic response, involving several genes in different chromosomes. Drought tolerance may be achieved through different mechanism ranging from morphological/ structural changes to physiological and biochemical processes such as osmotic adjustment, activation of antioxidant defences and hormone synthesis (Xie et al., 2023). The predominant phytohormone involved in drought response is ABA (Wei et al., 2025). During plant responses to drought, two major groups of genes are involved. The first group comprises genes encoding structural proteins, which function as downstream effectors in the stress response pathway. These include osmo-regulatory genes, antioxidant proteins, aquaporins, and late embryogenesis proteins (Bailly et al., 2001; Breton et al., 2003; Shinozaki & Yamaguchi-Shinozaki, 2007; Wang et al., 2006). The second group consists of genes encoding regulatory proteins, which act as early response transcription activators. These include transcription factors (bZIP, WRKY, MYB, AP2/EREBP) and protein kinases, which shown to play a crucial role in the regulation of drought tolerance (Kim et al., 2024). Therefore, our findings show that GWAS revealed fifteen genomic regions across eight chromosomes associated with drought response. These associations involved interactions between drought and multiple traits at different developmental stages. Altogether, the results highlight the potential involvement of distinct drought response mechanisms in this population.

Twenty-two candidate genes associated with drought response were identified, spanning biochemical, physiological, and morphological traits according to their annotated functions. Regarding earliness traits, the response mechanism can be drought avoidance in the case of delayed flowering or drought escape in the case of earlier flowering. The overall trend in the population was a delayed flowering. Five genomic regions in chromosomes 2, 3, 7 and 12 were associated with earliness traits in different stages (Table S2). In chromosome 2, U2AF2 (Solyc02g069790) was identified as associated with earliness. In *Arabidopsis* U2AF65 (U2AF2) regulates flowering time and pollen-tube growth, and the *AtU2AF65A* mutant exhibits late flowering(Park et al., 2019). *AtU2AF65A* plays a role in ABA-mediated splicing regulation, influencing flowering time (Xiong et al., 2019). Since ABA is a key regulator of drought response, this suggests that U2AF2 may be a flowering time regulator under drought conditions.

Three genomic regions associated with drought response in vegetative traits were found. In chromosome 1, SNF4-like protein (Solyc01g099280) was identified as a candidate gene related with plant height. A study on transgenic lines of TaSnRK2.4, a member of the SNF gene family in wheat, showed that these plants exhibited lower leaf water loss upon detachment, higher root water content, and increased osmotic potential in seedlings compared to controls. These plants also demonstrated more robust photosynthetic performance under moderate drought stress, suggesting that TaSnRK2.4 confers improved drought resilience (Mao et al., 2010). Regarding total biomass production, the three suggested candidate genes malate synthase (Solyc03g111150), 4CL (Solyc03g111170) and adenylate kinase (Solyc03g111200) in chromosome 3 have different functions that may affect overall biomass production under drought conditions. Therefore, all three genes may be involved in total biomass production in WS, as their functions range from energy transport and osmotic regulation to lignin production and flavonoid synthesis (Gholizadeh et al., 2025; Peracchi et al., 2024; Rajeev et al., 2025).

The candidate gene detected in chromosome 2, associated with wilting2, is a Late embryogenesis abundant (LEA) (Solyc02g078930), which is known to participate in plant stress tolerance response. The primary physical characteristic of LEA proteins is that they are typically hydrophilic. Under drought conditions, LEA proteins are rapidly upregulated to maintain membrane stability and reduce water loss. Increasing the expression of LEA genes in rice, such as HVA1 and LEA3-1, delays leaf wilting and prevents premature leaf death (Yu et al., 2016). In terms of leaf loss, it is characterized by cellular degradation often triggered by environmental factors such as drought or age, leading to leaf yellowing and eventual abscission. The suggested candidate gene DCD (Development and Cell Death) domain protein (Solyc09g074380) is an orthologue of At5G42050 and At3G27090, two development and cell death domain-containing asparagine-rich protein (DCD/NRP) that can induce cell death when transiently expressed. A delay in developmentally programmed leaf senescence was observed in association with these proteins (Camargos et al., 2019).

Proline, acts as an osmoprotectant and plays a role in scavenging reactive oxygen species (ROS), thereby protecting plants from oxidative damage, which is considered a drought tolerance mechanism(Per et al., 2017). C2H2 zing finger gene (Solyc07g063970) detected as a candidate gene in chromosome 7, is a component of the ABA pathway, and increases stress tolerance by acting in other pathways. In sweet potatoes, overexpression of IbZFP1, another C2H2 zinc finger protein, significantly upregulated genes related to ABA signalling, proline biosynthesis, and ROS scavenging (Wang et al., 2016; Wang et al., 2019). For stomatal water conductance (g_sw_), which is the movement of water vapor through the stomata, plants often close their stomata to reduce water loss as a drought avoidance mechanism. Different genomic regions identified for g_ws_1 and g_ws_2 indicate different genetic control regulating stomatal conductance at different developmental stages. The candidate gene Abscisic Acid Receptor PYL10 (Solyc01g095700), associated with g_ws_2, is directly involved in regulating stomatal conductance. PYR/PYL receptors play a crucial role in basal ABA signalling, which is essential for regulating stomatal aperture, and progressive inactivation of PYR/PYL genes results in an additive increase in g_sw_ (Wei et al., 2025).

In summary, different candidate genes located in distinct genomic regions were linked to flowering time and stomatal regulation under drought conditions, with their involvement varying based on the developmental phase, shedding light on the wide genetic control of drought stress response that varies depending on the trait type and the mechanisms plants employ to cope with drought, with a strong dependence on the developmental stage. In some cases, multiple candidate genes were identified within the same genomic region based on their functional roles, which may suggest a polygenic basis for these traits or the existence of a drought-associated hotspot.

## 5. Conclusions

Findings of this study, including results from genomic assisted selection and GWAS, highlight the complexity and polygenic nature of drought response mechanisms in plants. On one hand, the analysis underscores the involvement of multiple genetic regions, each contributing to various aspects of drought response. On the other hand, it emphasizes the significance of the ToMAGIC population in uncovering these genetic regions, which were shaped by extensive recombination events during the population’s development. These recombination events led to the formation of transgressive lines, which exhibit superior traits beyond the parental phenotypic range, making them highly valuable for improving tomato performance under drought conditions. Consequently, these transgressive lines offer a unique genetic resource that can be directly integrated into breeding programs, either as rootstocks to enhance drought resilience in grafted plants or as pre-breeding materials to improve drought adaptation in future cultivars. Their incorporation into breeding pipelines can accelerate the development of tomato crops with enhanced performance under water-limited conditions, benefiting both agricultural productivity and environmental sustainability.

## Supplementary tables

Table S1. List of traits evaluated.

Table S2. Pairwise Pearson correlations within each treatment (C and WS) and their respective significance (p value).

Table S3. GWAS results highlighting significant SNPs exceeding the Bonferroni threshold and candidate genes in each associated region

---

**Figure S1.**
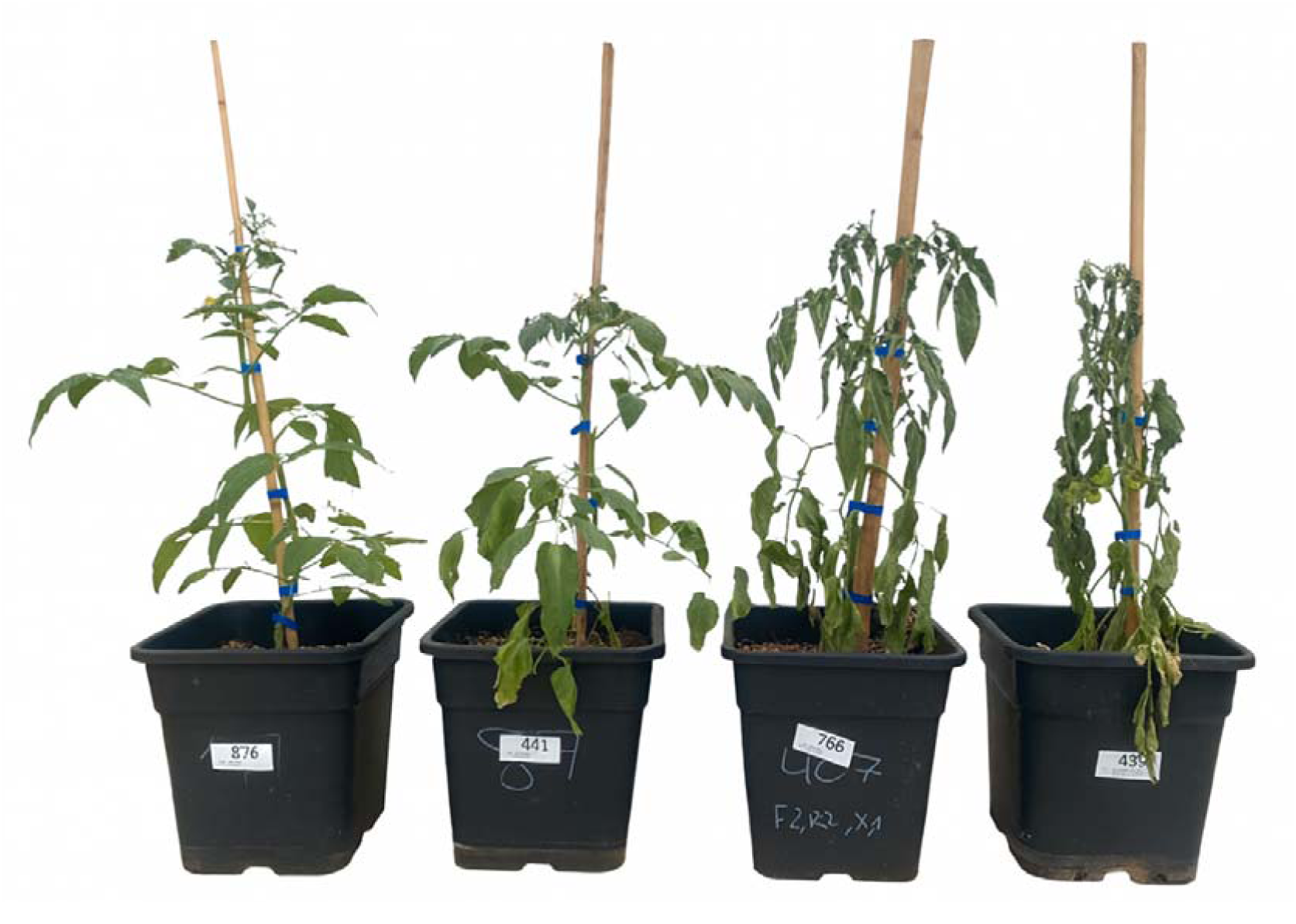
Wilting scale used to evaluate the effect of water stress (WS) on plant turgor. From left to right: 1 = <25 % of leaves wilted; 2 = 25–50 % wilted; 3 = 50–80 % wilted; 4 = severe wilting across the entire plant. Measurements were taken at 48–50 DAT and 80–82 DAT.

**Figure S2.**
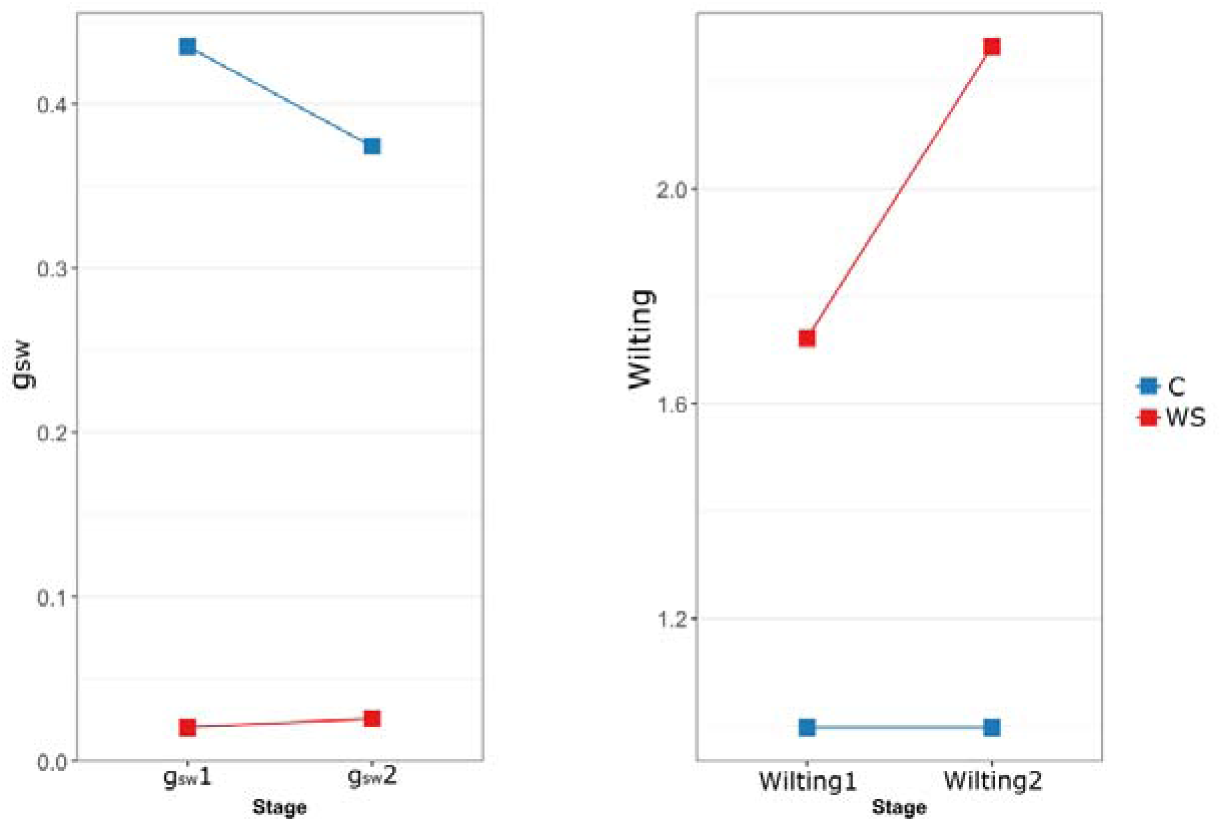
Sequential measurements of g_sw_ and wilting under water stress (WS, red line) and control (C, blue line) conditions. Plots display the estimated marginal means for the CCToMAGIC population after each WS period.

